# A protein truncating R179X variant in *RNF186* confers protection against ulcerative colitis

**DOI:** 10.1101/035105

**Authors:** MA Rivas, D Graham, P Sulem, C Stevens, AN Desch, P Goyette, D Gudbjartsson, I Jonsdottir, U Thorsteinsdottir, F Degenhardt, S Mucha, MI Kurki, D Li, M D’Amato, V Annese, S Vermeire, R Weersma, J Halfvarson, P Paavola-Sakki, M Lappalainen, M Lek, B Cummings, T Tukianen, T Haritunians, L Halme, LLE Koskinen, A Ananthakrishnan, Y Luo, GA Heap, M Visschedijk, NIDDK IBD Genetics Consortium, UK IBD Genetics Consortium, DG MacArthur, BM Neale, T Ahmad, CA Anderson, SR Brant, R Duerr, M Silverberg, J Cho, A Palotie, P Saavalainen, K Kontula, M Färkkilä, DPB McGovern, A Franke, K Stefansson, JD Rioux, RJ Xavier, MJ Daly

## Abstract

We conducted a search for protein truncating variants conferring protection against inflammatory bowel disease exploiting knowledge of common variants associated with the same disease. We found that a protein truncating variant (rs36095412, p.R179X, genotyped in 11,148 ulcerative colitis patients and 295,446 controls, MAF = up to 0.78%) in *RNF186*, a single-exon ring finger E3 ligase with strong colonic expression, protects against ulcerative colitis (overall *P* = 6.89×10^−7^, odds ratio (OR) = 0.30). We further demonstrate that the truncated protein is expressed, suggesting the protective mechanism may reside in the loss of an interaction or function via mislocalization or loss of an essential protein element.

## Main

### Motivation

A total of 200 loci have been unequivocally implicated to the two common forms of inflammatory bowel diseases (IBD): Crohn’s disease (CD) and ulcerative colitis (UC)^1,2^. For these findings, like most GWAS results, it has proven challenging to infer the functional consequences of common variant associations^3^ beyond cases where protein-altering variants have been directly implicated. Protein-truncating variants (PTVs), also commonly referred to as loss-of-function (LoF) variants^4^ as they often result in a non-functional or unstable gene product, are generally the strongest acting genetic variants in medical genetics and, as one functional copy of the gene is removed, may often provide insight into what is achievable pharmacologically via inhibition of the product of the gene^5^. Thus, identifying PTVs that are demonstrated to lead to loss of gene function and confer protection from disease hold particular promise for identifying therapeutic targets^6–9^.

### Screen Sequencing

We conducted targeted sequencing of the exons of 759 protein-coding genes in regions harboring common variants associated to IBD^10,11^ in 917 healthy controls, 887 individuals with UC (cases), and 1204 individuals with CD (cases) from the NIDDK IBD Genetics Consortium (North American clinical samples of European descent). We jointly analyzed these data with sequencing data from the same genes taken from an exome sequencing data set of Finnish individuals: 508 with UC, 238 with CD, and 8124 Finnish reference samples sequenced within Sequencing Initiative Suomi (SISu) project (www.sisuproject.fi)^12^. Across this targeted gene set, we discovered 81 PTVs found in 2 or more individuals (Supplementary Table 1) and used a Cochran-Mantel-Haenszel chi-squared test to scan for protective variants with two strata corresponding to the two cohorts. The test for association was run based on the phenotype (CD, UC or IBD) indicated by the common variant association in the region^13^ (that is, truncating variants in a gene associated only to CD such as at *NOD2* would be tested for CD vs. control association). We identified three putatively protective PTVs with a *P* value < 0.05: (1) a previously published low-frequency variant in *CARD9* (c.IVS11+1G>C) located on the donor site of exon 11 which disrupts splicing (*P* = 0.04)^7,14^; (2) a frameshift indel in *ABCA7* (*P* = 0.02); and (3) a premature stop gain variant (p.R179X) in *RNF186* with signal of association (*P* = 0.02) to ulcerative colitis. As the *CARD9* result was a well-established protective association, and *ABCA7* contained four additional PTVs that did not appear protective (combined odds ratio was greater than 1 and a signal of mixture of risk and protective effects would require a sequencing follow-up strategy), we focused specifically on follow-up work to confirm or refute the association of the *RNF186* nonsense variant (the only PTV detected in *RNF186* in either sequence data set).

**Table 1.**
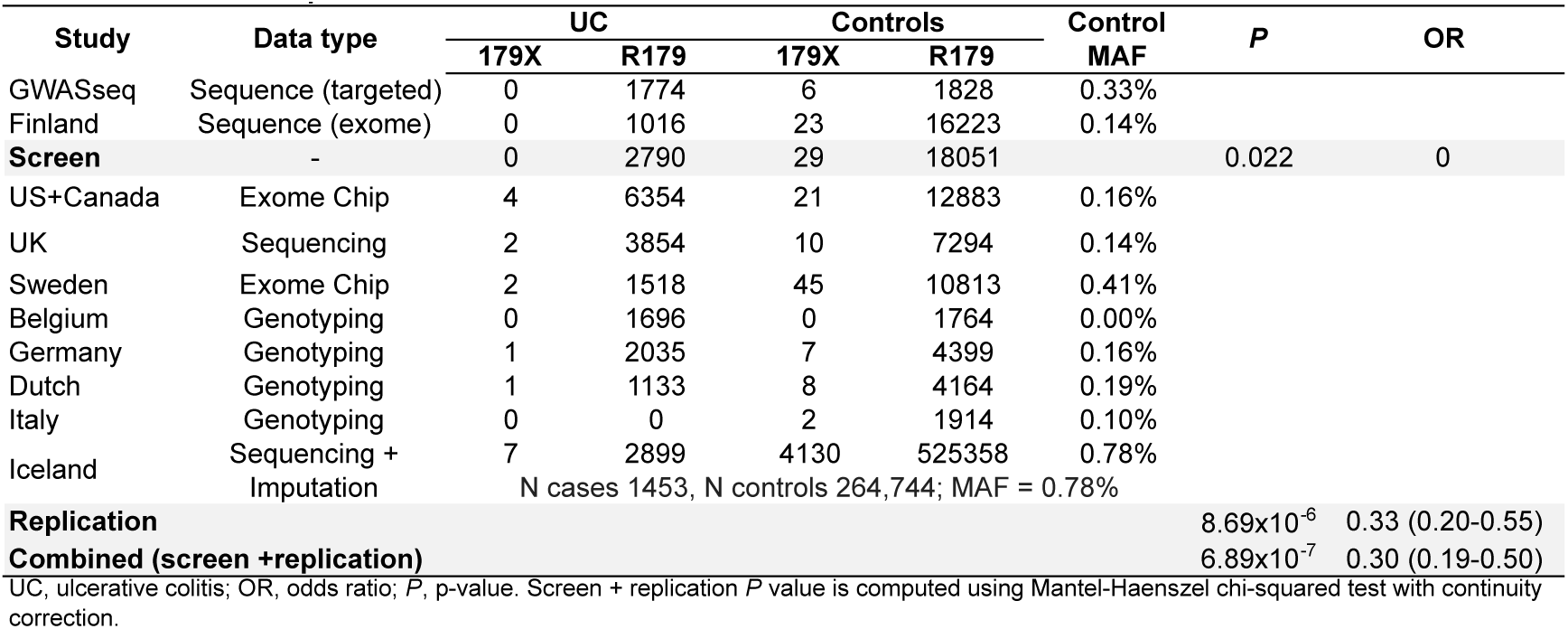
Association of p.R179X in *RNF186* with ulcerative colitis

### Replication

Replication genotype data obtained in 8,300 ulcerative colitis patients and 21,662 controls from US, Canada, UK, Sweden, Belgium, Germany, Netherlands and Italy provided strong support that the premature stop-gain allele, p.R179X, confers protection against ulcerative colitis (*P* = 0.0028, OR = 0.36 [95% CI=0.19-0.71]). Cluster plots from all genotyping assays were manually inspected to insure consistent high quality across all experimental modalities used to assess this variant (Supplementary figures 1,2).

Further evidence of replication was seen in whole-genome sequence data followed by imputation collected by deCODE Genetics^15,16^, in which a set of 1,453 Icelandic patients with UC were compared to a very large population sample (n=264,744) and a consistent strong protection (*P* = 5.0×10^−4^, OR = 0.30 [95% CI=0.15-0.59], imputation information of 0.99; overall replication *P* = 8.69×10^−6^, OR = 0.33 [0.20-0.55]) was observed between the truncating allele and the disease (Table 1, Materials and Methods). Of note, this observation is advantaged by the property that R179X has a roughly four-fold higher frequency in Iceland (MAF=0.78%) than in other European populations such that the Icelandic group, despite a moderate contribution in absolute number of cases (close to 1/6), have around half of the contribution in term of effective sample size and power.

In Iceland, the sequencing of a large fraction of the population followed by imputation assisted by long range phasing in around 150 thousands individuals allow us to identify homozygotes for rare loss-of-function variants^17^. We find 8 individuals homozygous for the 179X allele (MAF=0.78%), the oldest reached the age of 70 (still alive) and one of the eight died (age 62), consistent with Hardy-Weinberg expectation (n=9.1). There was no significant association of the homozygous genotype with a decreased lifespan or fertility (number of children). Given the lower frequency in other populations, no homozygous individual was expected and/or detected in the Exome Aggregation Consortium (ExAC) nor in the remaining set of individuals in this study. Together this indicates that having 2 copies with a stop gained in RNF186 is compatible with life, reproduction and aging – and importantly does not highlight severe medical consequences that would be of obvious concern in developing a therapeutic to mimic the effect of this allele.

The combined significance across all samples tested is *P* = 6.89×10^−7^ (OR = 0.30 [95% CI=0.19-0.50]) – considering we advanced only one variant to follow-up study, the replication p-value of 8.7×10^−6^ is unequivocally significant and would have been significant even if 5000 variants were put through this follow-up, let alone all 81 that were discovered in the sequencing screen. No other PTV in *RNF186* was discovered and tested in our sequencing. The gene is relatively small (227 amino acids) and R179X is by far the most common detected PTV in EXAC, more than 10 times more common than the sum of all other PTVs in the gene.

In the initial UC GWAS study the 1p36 locus had at least two non-coding independent association signals separated by recombination hotspots (rs4654903 and rs3806308; r^2^ = .001, minor allele frequency = 45.5% and 47%, respectively,) that did not implicate any one of the three genes (RNF186-OTUD3-PLA2G2E) in the region^18^. Recently, a low-frequency coding variant in *RNF186* (rs41264113, p.A64T, MAF=0.8%) was found to confer increased risk to ulcerative colitis (OR = 1.49 [1.17-1.90])^14^. In the discovery and replication component of this study p.R179X was found to lie on the haplotype background of the risk allele for rs4654903 and very little correlation was observed with rs3806308 or the low-frequency coding variant p.A64T (Supplementary figure 3). Naturally, the protective signal at p.R179X remains unchanged when corrected for the background higher risk allele rs4654903 on the haplotype it arose on (Supplementary table 2).

### Profile of expression of RNF186 p.R179X: transcript and protein expression

*RNF186*, a single-exon protein-coding gene, encodes the ring finger E3 ligase which localizes at the endoplasmic reticulum (ER) and regulates ER stress-mediated apoptosis in a caspase-dependent manner^19^. To understand the functional consequences of p.R179X we integrated transcriptome and protein expression level data. First, we examined the gene expression profile of *RNF186* (encoded by a single transcript isoform: ENST00000375121) across multiple tissues in the Genotype Tissue Expression (GTEx) project and identify that its highest expression is in the transverse colon (median RPKM = 17.32, *n* = 61) with only three other tissues having an RPKM level above 1: i) pancreas, ii) kidney cortex, and iii) the terminal ileum (Supplementary figure 4A)^20^. RNF186 protein was observed at “medium” expression levels for tissues in the gastrointestinal tract (stomach, duodenum, small intestine, appendix, colon) in the Human Protein Atlas^21^ (Supplementary figure 4B).

### Functional consequences of RNF186 p.R179X: impact on transcript allele expression

We integrated allele-specific expression (ASE) data for individual carriers of p.R179X in the GTEx project. Within GTEx we identified, two individual carriers with some containing ASE data for p.R179X in transverse colon and sigmoid colon. The carriers had consistent patterns of no ASE effects (Supplementary figure 5,6) suggesting that nonsense-mediated decay (NMD) does not degrade the aberrant transcripts containing the truncating alleles and that additional functional follow-up would be necessary to determine the molecular impact of p.R179X on RNF186^5^. The gene contains one exon and is intronless, and there is prior expectation that those genes do not undergo nonsense mediated decay since this presence is reported to requite the presence of at least one intron.

### Functional consequences of RNF186 p.R179X: impact on protein allele expression

Given that R179X messenger RNA was detected at levels similar to the common allele, we sought to quantify protein expression. Accordingly, we transiently transfected 293T cells with RNF186 expression constructs containing an epitope tag for detection on western blot with anti-V5 antibodies. As expected, we found that the common allele of RNF186 was efficiently expressed at the protein level, and strikingly, so was R179X (Fig. 1). Although we detected expression of RNF186 regardless of the position of the V5 tag, R179X protein was detected only when the V5 tag was present on its N-terminus (Fig. 1). Notably, the common allele encodes two transmembrane domains and lacks an N-terminal signal peptide, which supports a model in which RNF186 N- and C-termini are present on the cytoplasmic side of membrane structures. In contrast, R179X lacks the second transmembrane domain, and must therefore position its N- and C-termini on opposite sides of the membrane. Thus, the charged C-terminal V5 tag appended to R179X cannot efficiently translocate across the ER membrane into the lumen during translation. In contrast, the N-terminal V5 tag did not impair expression of R179X, likely because it is present on the cytoplasmic side of the membrane and does not require membrane translocation. Collectively, we conclude that R179X truncation does not preclude protein expression.

**Figure 1.**
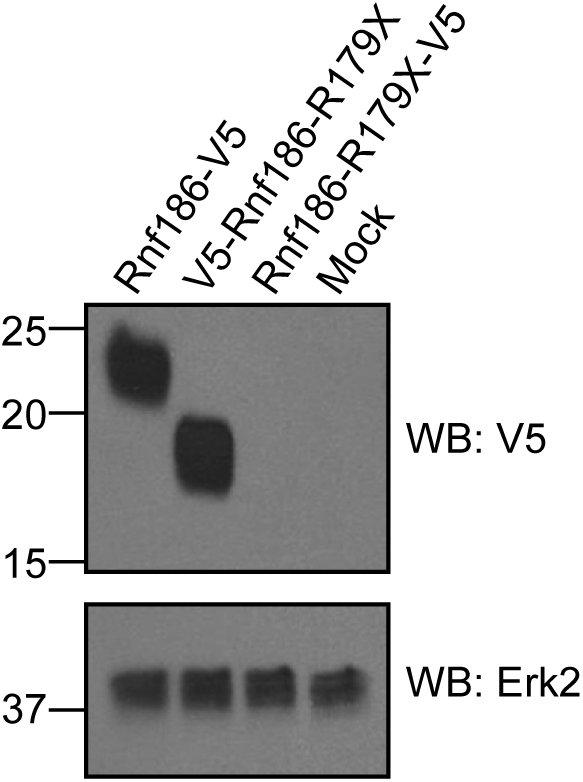
A. 293T cells were transfected with the indicated expression constructs and analyzed by western blot for expression of Rnf186 and the R179X variant. Western blot for Erk2 was performed as a loading control. The RNF186 protein with a premature stop at amino acid position 179 is expressed. B. 293T cells were transfected with the indicated expression constructs and analysed by immunofluorescence imaging for subcellular localization of Rnf186 and the R179X variant.

This study strengthens the direct evidence for the involvement of *RNF186* in ulcerative colitis risk in which a powerful allelic series, including common non-coding alleles and risk increasing p.A64T is available for further experimentation. Further supporting the medical relevance of this truncating variant, the same rare allele of the same variant coding p.R179X has been reported in Iceland to have a genome wide significant association with a modest increase in serum creatinine level (effect = 0.13 SD, *P* = 5.7 × 10^−10^) and a modest increase in risk of chronic kidney disease^16^. In the context of the few other established protective variants in IBD, including the coding *IL23R* variants (p.V362I, MAF = 1.27%, OR = 0.72 [0.63-0.83]; p.G149R, MAF = 0.45%, OR = 0.60 [0.45-.0.79]) and the splice disrupting *CARD9* variant (c.IVS11+1G>C, MAF = 0.58%, OR = 0.29 [0.22-0.37])^7^, R179X exerts a comparable protective effect. Because of the strong protective effect associated with the *RNF186* protein truncating variant, studies of RNF186 inhibition and the specific action of this variant protein should be useful in understanding the mechanism by which protection to ulcerative colitis disease occurs and whether this reveals a promising therapeutic opportunity similar to that which has been realized from the example of PCSK9 and cardiovascular disease.

## Materials and Methods

### Cohort descriptions

For all cohorts, UC was diagnosed according to accepted clinical, endoscopic, radiological and histological findings.

Genotyping of the Belgian cohort was performed at the Laboratory for Genetics and Genomic Medicine of Inflammatory (www.medgeni.org) of the Université de Montréal. Belgian patients were all recruited at the IBD unit of the University Hospital Leuven, Belgium; control samples are all unrelated, and without family history of IBD or other immune related disorders.

NIDDK IBD Genetics Consortium (IBDGC) samples were recruited by the centers included in the NIDDK IBDGC: Cedars Sinai, Johns Hopkins University, University of Chicago and Yale, University of Montreal, University of Pittsburgh and University of Toronto. Additional samples were obtained from the Queensland Institute for Medical Research, Emory University and the University of Utah. Medical history was collected with standardized NIDDK IBDGC phenotype forms. Healthy controls are defined as those with no personal or family history of IBD. The Prospective Registry in IBD Study at MGH (PRISM) is a referral center-based, prospective cohort of IBD patients. Enrollment began 1 January 2005. PRISM research protocols were reviewed and approved by the Partners Human Research Committee (#2004-P-001067), and all experiments adhered to the regulations of this review board. The PRISM study data were merged with population controls of European ancestry broadly consented for biomedical studies. These controls included samples from the NIMH repository^22^, POPRES^23^, the 1000 Genomes Project^24^ and controls ascertained for an age-related macular degeneration study^25^. The Italian samples were collected at the S. Giovanni Rotondo “CSS” (SGRC) Hospital in Italy. The Dutch cohort is composed of ulcerative colitis cases recruited through the Inflammatory Bowel Disease unit of the University Medical Center Groningen (Groningen), the Academic Medical Center (Amsterdam), the Leiden University Medical Center (Leiden) and the Radboud University Medical Center (Nijmegen), and of healthy controls of self-declared European ancestry from volunteers at the University Medical Center (Utrecht).

Subject ascertainment, diagnosis and validation for the UK samples are described elsewhere and are part of the UK Inflammatory Bowel Disease Genetics Consortium (UKIBDGC)^26^.

German patients were recruited either at the Department of General Internal Medicine of the Christian-Albrechts-University Kiel, the Charité University Hospital Berlin, through local outpatient services, or nationwide with the support of the German Crohn and Colitis Foundation. German healthy control individuals were obtained from the popgen biobank. Genotyping of the German cohort was performed at the Institute for Clinical Molecular Biology.

Finnish patients were recruited from Helsinki University Hospital and described in more detail previously^27,28^.

Subject ascertainment, diagnosis and validation for a subset of the Swedish samples with ulcerative colitis^14^ and without ulcerative colitis^29^ are described elsewhere.

Icelandic population: A total of 1,453 individuals diagnosed with ulcerative colitis, was used in the analysis. All the cases were histologically verified, and diagnosed either by 1997 or prospectively during the period 1997 to 2009 at Landspitali, the National University Hospital of Iceland.

### NIDDK NHGRI targeted sequencing

**Sample selection**. We selected 3,008 samples (1,204 crohn’s disease; 887 ulcerative colitis; and 917 controls) for sequencing composed of North-American samples of European descent from the NIDDK IBD Genetics Consortium repository samples.

**Target selection**. Target exonic sequences were selected based on the coding exons of 759 genes (2.546 megabases). Genes were selected if they were in regions identified in the genome-wide association studies for inflammatory bowel disease in Franke et al. 2010 (CD)^10^, Anderson et al. 2011 (UC)^11^.

### Finland exome sequencing

Finnish individuals were exome sequenced as part of Sequencing Initiative SUomi (SISu, www.sisuproject.fi). The SISu project consists of the following population and case control cohorts: 1000 Genomes Project, ADGEN (Genetic, epigenetic and molecular identification of novel Alzheimer’s disease related genes and pathways) Study, The Botnia (Diabetes in Western Finland) Study, EUFAM (European Study of Familial Dyslipidemias), The national FINRISK Study, FUSION (Finland-United States Investigation of NIDDM Genetics) Study, Health 2000 Survey, Inflammatory Bowel Disease Study, METSIM (METabolic Syndrome In Men) Study, Migraine Family Study, Oulu Dyslipidemia Families, Northern Finland Intellectual Disability (NFID) and Northern Finland Birth Cohort (NFBC). All samples were sequenced at the Broad Institute of MIT and Harvard, Cambridge US, University of Washington in St. Louis, US and Wellcome Trust Sanger Institute, Cambridge, UK.

**Exome sequencing**. To produce harmonized good quality call set we applied the core variant calling workflow for exome sequencing data that is composed of two stages that are performed sequentially: Pre-processing: from raw sequence reads to analysis-ready reads; and Variant discovery: from analysis-ready reads to analysis-ready variants (Supplementary Methods).

**Identification of Finnish samples**. In order to obtain genetically well-matched controls for comparison with the Finnish IBD cases we first jointly called Finnish exomes with Swedish exomes to identify genetic Finns from other close Nordic country. Final probability was obtained by dividing the probability of being Finnish divided by the sum of probabilities of being Finnish or Swedish. Training samples in distance calculations were selected for being from Finnish or Swedish cohort as appropriate and clustering on the expected cluster (PC1 < 0.002 for Finnish samples and PC1 > =0 and PC2<=0.01). Samples with > = 99% Finnish probability (8124 non-IBD samples; 508 UC; 238 CD; 92 IC) were then subset and PCA with the same parameters was run again to obtain PCA’s for Finnish substructure (Supplementary Material).

### Variant annotation

Variants for the targeted and exome sequencing datasets were annotated using PLINK/SEQ v0.10 and RefSeq reference transcript set downloaded from https://atgu.mgh.harvard.edu/plinkseq/resources.shtml.

### Follow-up genotyping of *RNF186*

**Sequenom**. *RNF186* p.R179X was assayed using Sequenom MassARRAY iPLEX GOLD chemistry and SpectroCHIPs were analyzed in automated mode by a MassArray MALDI-TOF Compact system 2 with a solid phase laser mass spectrometer (Bruker Daltonics Inc.). The variant was called by real-time SpectroCaller algorithm, analyzed by SpectroTyper v.4.0 software and clusters were manually reviewed for validation of genotype calls. Reported genetic map positions for the markers were retrieved from the SNP database of the National Center for Biotechnology Information (NCBI).

**Exome array**. The Illumina HumanExome Beadchip array includes 247,870 markers focused on protein-altering variants selected from >12,000 exome and genome sequences representing multiple ethnicities and complex traits. Nonsynonymous variants had to be observed three or more times in at least two studies, splicing and stop-altering variants two or more times in at least two studies. Additional array content includes variants associated with complex traits in previous GWAS, HLA tags, ancestry informative markers, markers for identity-by-descent estimation, and random synonymous SNPs. We focused on variant exm26442, which was the only PTV in the targeted sequencing data set that was also in the exome array and had a *P* value < 0.05 in the screening component of the study. Samples in the targeted sequencing data set were excluded from the exome array analysis.

**UK sequencing**. The UKIBDGC sequenced low coverage whole genomes of 1767 UC patients from our nationwide cohort (median depth 2X) and compared them to 3652 population controls from the UK10K project (median depth 7X). Samples were jointly called using samtools^30^, and subjected to two rounds of genotype improvement using BEAGLE^31^. Genotype count for R179X from exome sequencing data in 161 additional UK ulcerative colitis patients with severe adverse drug response to common IBD drugs were included.

### Replication in Iceland population

The Iceland population data has been extended following the step below^15,32^.

Sequencing was performed using three different types of Illumina sequencing instruments.

a. Standard TruSeq DNA library preparation method. Illumina GAIIx and/or HiSeq 2000 sequencers (n=*5582)*.
b. TruSeq DNA PCR-free library preparation method. Illumina HiSeq 2500 sequencers (n=2315).
c. TruSeq Nano DNA library preparation method. Illumina HiSeq X sequencers (n=556).

Genotyping and imputation methods and the association analysis method in the Icelandic samples were essentially as previously described^15^ with some modifications that are described here. In short, we sequenced the whole genomes of 8,453 Icelanders using Illumina technology to a mean depth of at least 10X (median 32X). SNPs and indels were identified and their genotypes determined using joint calling with the Genome Analysis Toolkit HaplotypeCaller (GATK version 3.3.0)^33^. Genotype calls were improved by using information about haplotype sharing, taking advantage of the fact that all the sequenced individuals had also been chip-typed and long range phased. The sequence variants identified in the 8,453 sequenced Icelanders were imputed into 150,656 Icelanders who had been genotyped with various Illumina SNP chips and their genotypes phased using long-range phasing^34,35^.

**Allele specific expression data for R179X carriers**. The primary and processed data used to generate the ASE analyses presented in this manuscript are available in the following locations: All primary sequence and clinical data files, and any other protected data, are deposited in and available from the database of Genotypes and Phenotypes (www.ncbi.nlm.nih.gov/gap) (phs000424.v6.p1). Tissues with at least eight reads of data are presented.

### Rnf186 protein expression and localization for R179X carriers

**Plasmids**. cDNA encoding human RNF186 was obtained from The Genetic Perturbation Platform (GPP, Broad Institute) and cloned by Gibson assembly into the pLX_TRC307 expression construct. Sequences encoding V5 and Flag tags were appended to oligonucleotides for PCR amplification of RNF186.

**Biochemistry**. 293T cells (ATCC) were transfected with RNF186 expression constructs by means of Lipofectamine 2000 (Life Technologies) as indicated by the manufacturer. One day after transfection, cells were lysed (1% NP-40 in PBS), resolved by SDS-PAGE, and detected by western blot. Mouse anti-V5 HRP (Sigma V2260-1VL) was diluted 1:5000 and used in conjunction with chemiluminescent substrate (Pierce SuperSignal West Pico).

### Ethics statement

All patients and control subjects provided informed consent. Recruitment protocols and consent forms were approved by Institutional Review Boards at each participating institutions. All DNA samples and data in this study were denominalized.

### Association analysis

Association analysis of protein truncating variants in targeted sequencing data and the exome sequencing data was performed using the Cochran-Mantel-Haenszel (CMH) chi-squared test implemented in R to screen for PTVs with evidence of protective signal of association^36^. Combined (screen + replication) association analysis was conducted with the CMH chi-squared test. In the replication cohort a set of 1,453 Icelandic patients with Ulcerative Colitis (UC) were compared to a very large group representing the general population (n=264,744). Logistic regression analysis was applied to the data set to obtain study specific association statistics.

**Online resources:** http://www.gtexportal.org/

## Acknowledgements

MJD is supported by grants from: the National Institute of Diabetes and Digestive and Kidney Disease (NIDDK) and the National Human Genome Research Institute (NHGRI; DK043351, DK064869, HG005923); the Crohns and Colitis Foundation (3765); the Leona M. & Harry B. Helmsley Charitable Trust (2015PG-IBD001); and Amgen (2013583217), RJX is supported by grants from Amgen (2013583217) and CCFA (3765). JDR is funded by grants from (DK064869; DK062432). IBD Research at Cedars-Sinai is supported by grant PO1DK046763 and the Cedars-Sinai F. Widjaja Foundation Inflammatory Bowel and Immunobiology Research Institute Research Funds. D.P.B.M. is supported by DK062413, AI067068 and U54DE023789-01, grant 305479 from the European Union, and The Leona M. and Harry B. Helmsley Charitable Trust and the Crohn’s and Colitis Foundation of America. SRB is support by an NIH U01 grant (DK062431).

